# The role of NMDA receptors in memory and prediction in cultured neural networks

**DOI:** 10.1101/2024.02.01.578348

**Authors:** Martina Lamberti, Michel J.A.M. van Putten, Sarah Marzen, Joost le Feber

**Author notes:** Martina Lamberti or Joost le Feber.

## Abstract

Memory has been extensively studied at the behavioural as well as the cellular level. Spike timing dependent plasticity (STDP) is associated with N-methyl-D-aspartate (NMDA) receptor activation and is widely accepted to be essential for long-term memory. However, experimental evidence remains sparse, probably due to the required complex combination of cellular and functional readouts. Recent work showed that in-vitro cortical networks memorize and predict inputs. The initial dependency of prediction on short-term memory decreased during the formation of long-term memory traces. Here, we stimulated in-vitro networks to investigate memory and prediction under control conditions, or under NMDA block. The NMDA anatagonist 2-amino-5-phosphonovaleric acid (APV) at a concentration that did not significantly reduce network excitability, but did impede long-term memory trace formation. In APV-treated cultures short-term memory of stimuli persisted, and they were still able to predict. In contrast to control cultures, prediction remained fully dependent on short-term memory. This confirms that NMDA receptor activation is essential for the formation of long-term memory traces and supports the notion that, as control cultures learn to memorize the stimulus, long-term memory starts to contribute to their predictive capability.

## Introduction

Memory is a crucial cognitive function for daily life [1, 2] that has been the subject of extensive investigation at various levels, ranging from cognitive psychological to neurobiological approaches. Cognitive psychology has mainly focused on abstract explanations of memory [3–5], and discriminates between short-term memory, acting on time scales of seconds, and long-term memory, on time scales of minutes to days [6–8]. New experiences are first stored in the hippocampus in a process called synaptic consolidation [9, 10]. Then, memories are transferred to the neocortex for long-term storage by repeated replay in the hippocampus, particularly during slow-wave sleep (system consolidation) [10, 11]. On the other hand, neurobiology has revealed detailed insights into neuronal functioning and synaptic plasticity that are likely to underlie memory [12]. Bridging the gap between cognitive and neurobiological studies has been challenging, at least in part because experimental paradigms usually do not facilitate assessment of parameters that cover the entire range from the molecular or cellular level to cognitive functioning.

It is widely accepted that plasticity of neuronal connectivity is crucial for memory. Connection strengths between neurons can be determined in patch clamp experiments [13], but this approach reveals only one, or at the best a few, connection strengths at a time. In past decades, micro electrode arrays (MEAs) have been used to record activity from networks of cultured neurons [14], and recorded activity patterns can be used to infer functional connectivity and study memory [15–17]. We will use the term ‘connectivity’ throughout the current work to refer to functional connectivity. Connectivity and activity in neuronal networks mutually affect each other. The finding that input deprived networks develop quasi stable activity patterns [18, 19] and connectivity [20] suggests that activity and connectivity are in equilibrium [21]. External stimuli induce new activity patterns, and disturb the equilibrium between activity and connectivity [22, 23]. After a while the network reaches a new equilibrium with spontaneous activity patterns that include the stimulus response, which is interpreted as the formation of long-term memory of the input. Consequently, repeated exposure to this stimulus induces no further connectivity changes. These observations suggest that memory traces are part of balanced connectivity and activity at the network level, and that detection of memory trace formation requires estimation of connectivity throughout networks. It is not feasible to determine network connectivity based on patch clamp recording of pre- and post synaptic signals, but MEA recordings do enable estimation of network connectivity. Recent work showed that cultured networks on MEAs are able to form memory traces when repeatedly exposed to a stimulus during a few hours, and that this ability depends on slow-wave sleep-like conditions. These preparations facilitate measurement and manipulation of cellular and synaptic properties, which provides the opportunity to bridge part of the gap between neurobiological and cognitive studies.

In addition to memory, prediction is also crucial to successfully conclude every-day actions. Prediction can be defined as the ability to reduce the uncertainty on upcoming external inputs, and is strongly related to memory [24–26]. In principle there can be no prediction without memory [25]. Most evidence to support the notion that neural networks are able to predict comes from theoretical work [1, 2, 24], and a few pioneering experimental studies using retinal [26–30] or hippocampal preparations [31–33].

Recent work also showed that random networks of dissociated cortical neurons are able to predict, suggesting that prediction is independent of specific network architecture [34]. In that study, prediction was shown to initially depend on short-term memory of given stimuli, which typically lasted several hundreds of milliseconds. During multiple hours of repeated focal stimulation, long-term memory traces were formed and prediction became less dependent on short-term memory. This suggests that focal stimulation-induced long-term memory traces also play a role in prediction. It remains unclear which mechanisms underlie prediction and to what extent long-term memory is involved.

Several studies have attributed the formation of long-term memory traces to spike timing dependent plasticity (STDP). [6, 35–42]. STDP governs changes in synaptic weights based on the timing of pre- and postsynaptic action potentials, and may lead to either long-term potentiation (LTP) or depression (LTD) of synaptic strengths [43–46]. Calcium influx mediated by NMDA receptor activation plays an important role in STDP [36, 43, 45, 47], and has been shown to affect the density of AMPA receptors in the synaptic region [36, 47, 48]. Depending on the amplitude and kinetics of the calcium influx, synapses can be either potentiated or depressed [36, 47, 49]. The strong association between memory and prediction suggests that STDP and NMDA receptor activation may also play a role in prediction.

In the current work, we present an in-vitro model to investigate memory and prediction. In particular we aimed to determine the involvement of NMDA receptor activation in memory and prediction. We applied focal electrical stimulation to networks of rat cortical neurons plated on MEAs, with and without pharmacological blockade of NMDA receptors, and investigated whether networks were able to i) form long-term memory traces, and ii) predict future stimuli. In addition, we determined to what extent prediction depended on (short-term or long-term) memory.

## Materials and methods

### Culture preparation

Cortical cells, obtained from newborn rats were dissociated by trypsin treatment and trituration, and then 60 µl droplets of cell suspension (about 50, 000 cells) were plated on multi electrode arrays (MEAs; Multi Channel Systems, Reutlingen, Germany), precoated with poly ethylene imine (PEI). The glass substrate contained 60 titanium nitride electrodes (diameter 30 µm and pitch 200 µm), on which a Plexiglas ring (diameter 20 mm) was glued to create a culture well (Figure 1A). The well was filled with 1 ml of Neurobasal commercial medium (Neurobasal-A medium -gluc -pyr, Thermo Fisher), supplemented with B27, 1.24 g/L glucose, Pen/strep/glutamine, 100 mg/L vitamin C and 10 ng/L NGF. MEAs were stored in an incubator, under standard conditions of 36^*◦*^C, high humidity, and 5% CO2 in air. The culture medium was refreshed twice a week by removing 300 µl and adding 400 µl of fresh medium, thus compensating for evaporation. All cultures were grown for at least 3 weeks before experiments started, to allow for culture maturation [20, 50, 51]. Mature cultures contained excitatory and inhibitory neurons [52], and astrocytes (ratio astrocytes neurons: 4:1) [53] (see Figure S3 in Supplementary Information). Before experiments, culture chambers were firmly sealed with watertight but O_2_ and CO_2_ permeable foil (MCS; ALA scientific). Then, MEAs were placed in a measurement setup outside the incubator. In this setup, standard conditions of 36^*◦*^C, high humidity, and 5% CO_2_, were maintained. Every recording began after a 15 minute accommodation period. At the end of each experiment, cultures were returned to the incubator. All surgical and experimental procedures were approved by the Dutch committee on animal use (Centrale Commissie Dierproeven; AVD110002016802), and complied with Dutch and European laws and guidelines. Results are presented in compliance with the ARRIVE guidelines.

**Fig 1.**
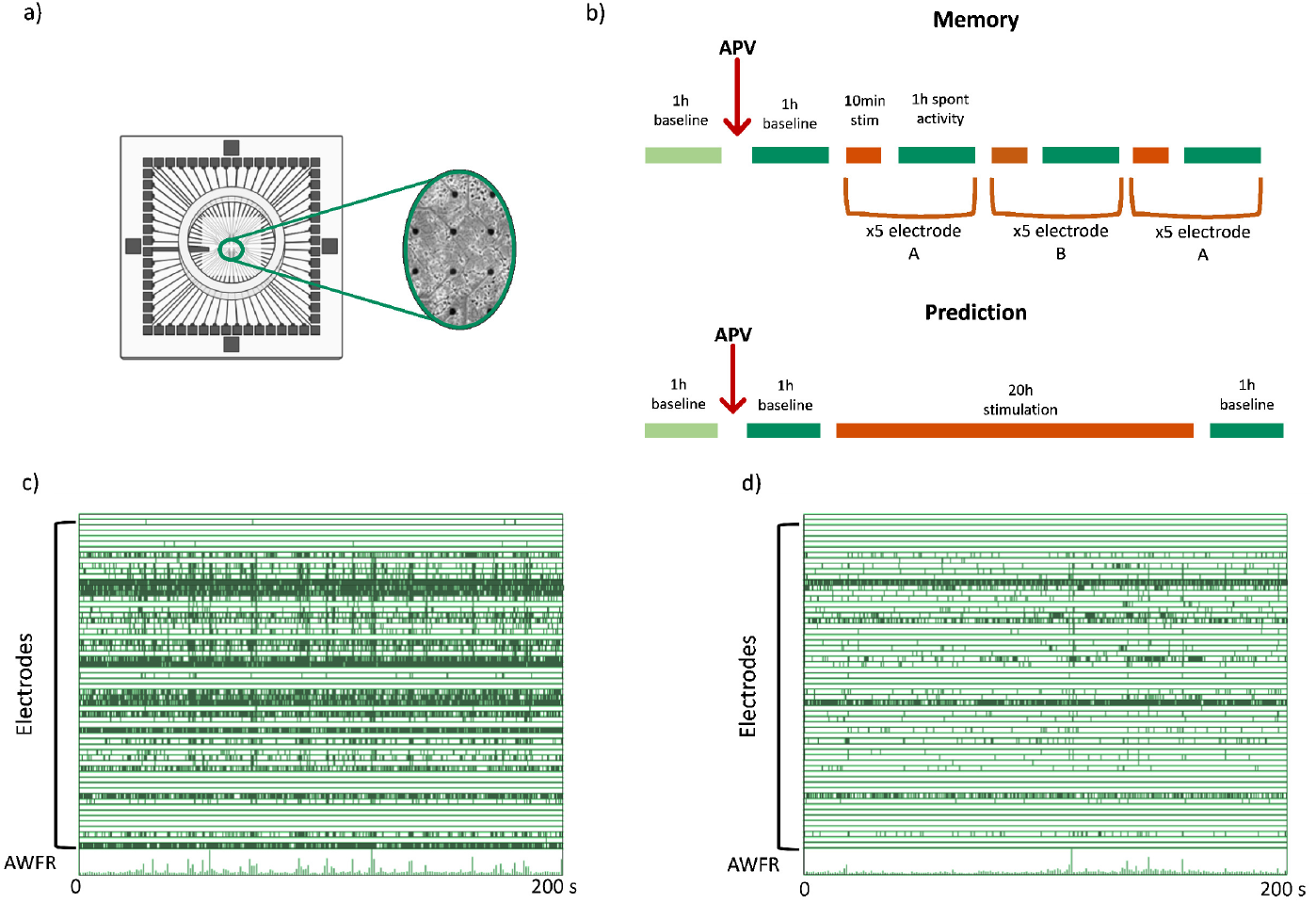
Example of MEA, experimental protocols and recorded activity. (a) shows an example of a 60 electrodes single well MEA used in this study. (b) shows the protocols for memory experiments (top) and prediction experiments (bottom). (c) shows an example of recorded spontaneous activity under control conditions. Each tick represents a recorded spike at the indicated electrode. Bottom trace shows summed activity (array wide firing rate; AWFR). (d) represents an example of spontaneous activity recorded after APV administration. (c) and (d) show that APV administration reduced the spontaneous firing rates, but firing patterns still contained synchronous activity (networks bursts)

### Recording set-up

To record activity, we placed MEAs into a setup outside the incubator, consisting of a MC1060BC preamplifier and FA60s filter amplifier (both MultiChannelSystems GmbH, Reut-lingen, Germany). Signals from the network were recorded by 59 electrodes using a custom-made Lab-View program, at a sampling frequency of 16 kHz per electrode.

Acquired analogue signals were band-pass filtered (2^*nd*^ order Butterworth 0.1 – 6 kHz) before sampling. Action potentials were detected whenever recorded voltages exceeded a threshold, set at 5.5 times the estimated root-mean-square noise level (ranging 3 – 5 µV). During recordings this noise estimation was continuously updated for each electrode. For each detected event the setup stores a time stamp, the electrode number, and the wave shape (6 ms). For off-line artifact detection and removal we used a wave shape based algorithm adapted from [54].

Focal stimulation was applied by electrical stimulation through one of the electrodes, using biphasic rectangular current pulses of 200 µs per phase [23]. After probing all electrodes at various amplitudes (16 *−* 24 µA), one electrode was selected for stimulation, using the lowest amplitude that induced a network response after at least 50% of all stimuli. Amplitudes were low enough to avoid electrolysis.

### Experimental Design

We used 20 mM 2-amino-5-phosphonovaleric acid (APV) to block NMDA receptors. This concentration clearly affected firing patterns and stimulation induced connectivity changes, but had no significant effect on network excitability (see Supplementary Information). Addition of APV was done under a laminar flow hood to maintain sterility. We performed experiments under control conditions (control cultures) or using an NMDA antagonist (APV cultures).

### Memory

We acquired 1 h of spontaneous activity at the beginning of the experiment (*Baseline*). After APV administration we recorded a second hour of spontaneous activity (*Baseline*_*AP V*_). Then both control and APV cultures underwent focal stimulation applied to two different electrodes (A and B). For each electrode we applied 5 stimulation periods of ten minutes each at a constant frequency of 0.2 Hz, alternated by 1 h spontaneous activity recordings. First, electrode A was used, then electrode B, and then again electrode A, leading to a total of 15 stimulation periods of 10 minutes. The total duration of the experimental protocol was 17 h and 30 min (Figure 1B).

### Prediction

To study prediction we adapted the protocol developed in earlier work [34]. In short, we first acquired 1 h of spontaneous activity under control conditions. After APV administration we recorded another hour of spontaneous activity. Then, cultures were subjected to 20 h of focal electrical stimulation (Figure 1B). For these experiments we used inter stimulus intervals (ISIs) drawn independently and identically from a density distribution designed to produce long-range temporal correlations, which were read from a pre-generated list (see Figure 4C). The long and intensive stimulation protocol adopted for the prediction experiments might be too stressful for the cultures to be applied twice. We therefore compared current results under NMDA blockade to control experiments from a previous study [34].

### Data analysis

#### Memory trace formation

Functional connectivity was inferred from spontaneous activity recordings using conditional firing probability (CFP) [20]. In short, CFP calculates the probability that electrode *j* spikes at time *t* = *τ*, given that electrode *i* spiked at *t* = 0. The maximum of this probability curve *M*_*i,j*_ was interpreted as the strength of the connection from electrode *i* to *j* (see Supplementary Information for details). For this analysis long-term recordings were divided into chunks that contained 2^13^ action potentials. This value was shown to be sufficient for statistical analysis of network connectivity in each chunk [23]. Still, with this chunk size, an hour of spontaneous activity could typically be divided into multiple chunks. This yielded a 60 x 60 connectivity matrix *M* for each chunk, that contained the strengths of connections between all possible pairs of electrodes. To assess the formation of memory traces, we quantified connectivity changes induced by each stimulation period by computation of Euclidean distances (EDs) between connectivity matrices [23] ( equation S2 in Supplementary Information).

Each stimulation period was preceded and followed by periods of spontaneous activity that contained multiple chunks of 2^13^ action potentials. Thus, we could in principle compute the distances between all possible pairs of baseline and post-stimulation chunks. However, this may not be necessary, if connectivity matrices hardly differ between baseline chunks, and if there was no significant drift. We calculated EDs between all possible pairs of chunks within baseline, and averaged them to determine the magnitude of spontaneous baseline connectivity fluctuations: *ED*_baseline_. To visualize possible drift, we calculated the Euclidean distances between the first baseline chunk and all other baseline chunks, and determined the slope (the average increase per unit of time) (see Figure 2). This analysis showed that the last chunk of baseline was representative of all chunks in baseline, and could be used to evaluate connectivity changes following electrical stimulation.

**Fig 2.**
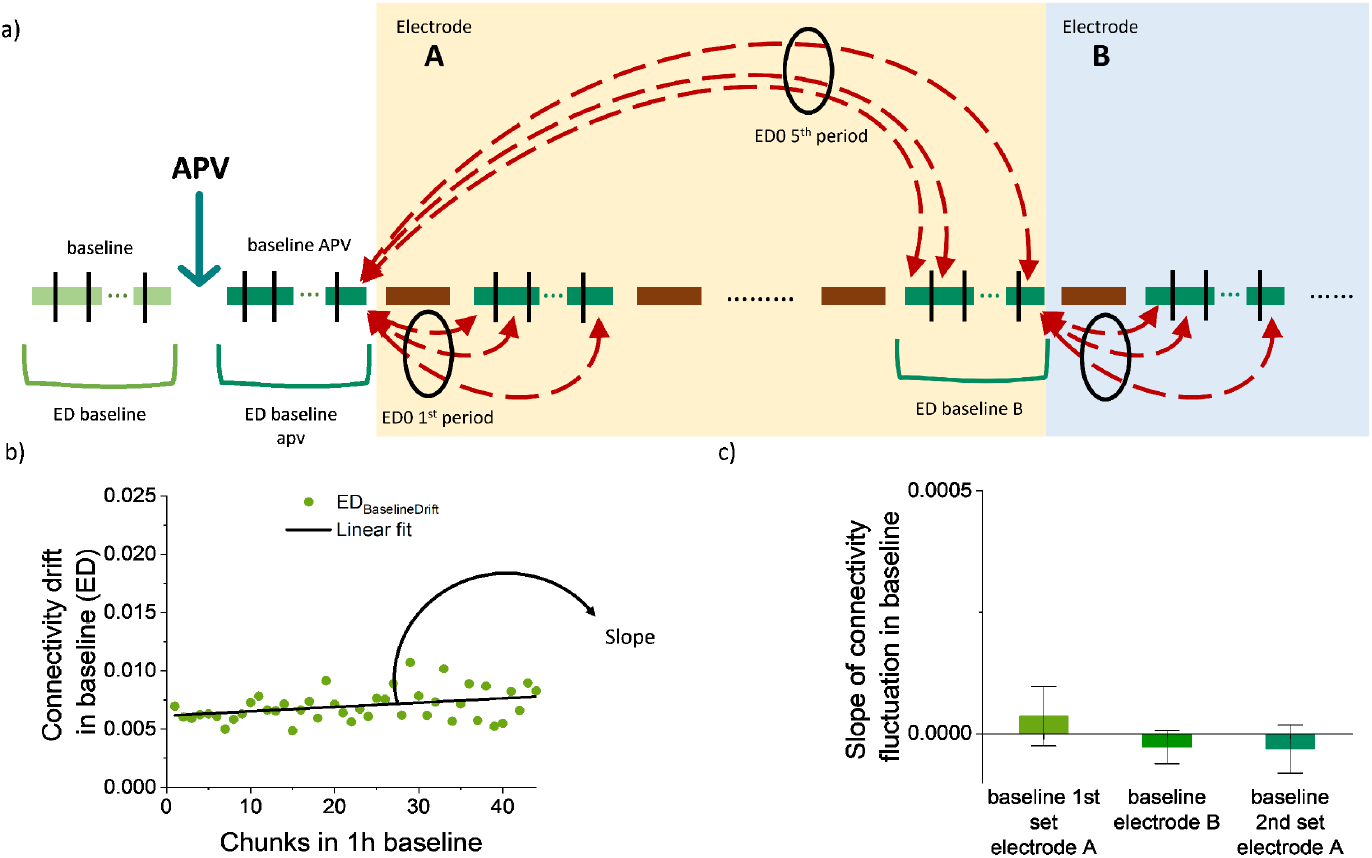
Calculation of Euclidean distances (ED). Top panel (a): visual representation of Euclidean distance computation in memory experiments. Black vertical lines indicates the division of spontaneous activity recordings into chunks, which were used to calculate connectivity matrices *M*. Red dashed arrows indicate Euclidean distance calculation between chunks and their corresponding baseline. Black circles indicate final averaging to obtain *ED*_0_ for each spontaneous activity period. Bottom panels indicate spontaneous connectivity drift in baseline recordings. Panel (b): Example of connectivity drift during baseline in one culture. Graph shows distances of all chunks from the first chunk. Drift was estimated as the slope of a linear fit, and averaged across cultures. Panel (c) shows mean drift during baseline, the last period of spontaneous activity before stimulation at electrode B, and the last one before return to electrode A. Error bars represents SEM.

Next, we computed EDs between the last chunk of baseline and each chunk contained in the (5) spontaneous periods that followed stimulation of electrode A. These distances are referred to as *ED*_0_. This procedure was repeated for electrode B, but now the last chunk before the first stimulation at electrode B was used as baseline. Similarly, after return to electrode A, the last chunk before the first (second round) stimulation at electrode A was used as baseline (see Figure 2). In the computation we considered only the nonzero elements of the *M* matrices. These analyses were applied to both controls and APV experimental data.

#### Network excitability

Network excitability, defined as the average network response to a spike in one neuron, was calculated with SPRs as developed in [55]. This allows estimation of the average response at electrode *j* to a single spike at electrode *i* under widely varying dynamic regimes. The computation of SPRs requires deconvolving the auto-correlation of electrode *i* from *CFP*_i,j_. Also in this case spontaneous recordings were subdivided in blocks of 2^13^ recorded spikes. SPR strengths of all pairs of active electrodes were calculated for each period of spontaneous activity in both control and APV groups, and averaged to obtain a measure of network excitability.

#### Quantification of prediction

We first verified that networks remained responsive to stimulation throughout experiments (see Supplementary Information). We used mutual information to quantify prediction [26, 34]. Mutual information estimates how much one signal reduces the uncertainty of another one. Then we first binarized the recorded neural activity (*X*) in bins of Δ*t* = 100 ms by setting *X*_*i*_ = 1 if electrode i recorded activity in that time bin, and *X*_*i*_ = 0 if not. The same binarization was applied to the stimulation signal (*S*). After binarizing the activity *X* and the stimulation signal *S*, for each hour, we calculated MI between *S* and *X*, with *X* shifted in time by *n* × Δ*t*, with −10 < *n* < 20. Negative shifts reveal memory, as quantified by *MI*_past_:

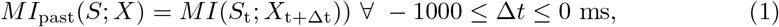

where *S*_t_ and *X*_t+Δt_ represent the unshifted stimulation signal and the time-shifted binarized activity. Activity was shifted forward up until a maximum of 2 seconds. Positive shifts are used to quantify prediction by computing *MI*_future_:

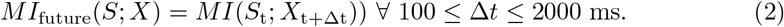

Consequently from Eqs. 1 and 2 *MI*_past_ and *MI*_future_ are functions of the shift Δ*t*. We applied this analysis to the APV prediction experiments and compared our results to control experiments obtained in earlier work [34]. In APV experiments, stimulus responses significantly decreased after 13 hours (see Supplementary Information, Fig S2). Therefore, we took into account only the first 10h of stimulation (instead of all 20h) for APV as well as control cultures.

#### Relationship between prediction and short-term memory

To study possible effects of APV on the relationship between short-term memory and prediction we analyzed the area under the curve of *MI*_future_ (Σ*MI*_future_) and *MI*_past_ (Σ*MI*_future_) for each analyzed hour, for each experiment (see Eq. S3 and S4 Supplementary Information) [34]. Then we plotted Σ*MI*_future_ against Σ*MI*_past_, and fitted a linear equation to determine the slope and offset of the relation, and the correlation coefficient. Finally, we investigated whether and how slope and offset changed during the considered 10 hours of stimulation.

#### Efficiency of predictions

We used the predictive information bottleneck concept as shown in Lamberti et al. [34] to determine whether APV affected the efficiency of prediction. This allows us to determine the minimum information about the past (*I*_mem_) necessary to make predictions on the future (*I*_pred_) (see Supplementary Information). The entire past and future were replaced by the time since the last stimulus (past; *S*^+^), and the time to the next stimulus (future; *S*^*−*^) [56]:

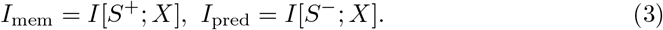

Prediction as quantified in experimental data under APV or control conditions was then compared to a theoretical optimum (*R*(*D*)). The theoretical optimum was obtained calculating the minimum memory required to enable a certain prediction. This resulted in a line that divides between achievable and unachievable combinations of information about the past (*I*_mem_) and the future (*I*_pred_).

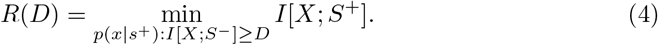

Here *D* corresponds to the required predictive power, *R*(*D*) is the memory necessary to reach that predictive power, *p*(*x*|*s*^+^) is the conditional probability of neural activity given time since the last stimulus, *x* is a realization of the activity *X, s*^+^ is a realization of *S*^+^. For both controls and APV treated cultures we computed *I*_mem_ and *I*_pred_ considering the first 10 h of stimulation only.

### Statistical Analysis

All analyses were performed using SPSS statistics for Windows (IBM, Inc., Chicago, IL) or Matlab (The Mathworks, Inc., Natick, MA, USA). Normality of data distribution was verified using Shapiro-Wilk tests. Effects of APV concentrations was testes with one sample t-test, in case of normality, or one sample Wilcoxon test otherwise.

Significance of connectivity changes was assessed using repeated measurements ANOVA, in case of normality, or repeated measurement Friedman test otherwise. Assessment of possible effects of APV on network excitability was performed with two way repeated measurement ANOVA, with condition and time as independent variables. In both memory and prediction experiments significance of temporal differences in effectiveness of stimulation were analyzed by one way ANOVA (if normally distributed) or Kruskal-Wallis (if non-normal). In the end t-statistics measure was applied to check trends of slope and offset of the relationship between memory and prediction. P-values

*<* 0.05 were considered to indicate significance.

## Results

### Long-term memory

In total 12 cultures were used, first under control conditions (control experiments), and then under NMDA blockade (APV experiments). All 12 control recordings were included for analysis, and 7 of the APV experiments. In the other 5 cultures APV administration reduced network activity to such low levels that CFP analysis could not be done to estimate functional connectivity.

### Network excitability

Network excitability was quantified by the average magnitude of SPRs during each period of spontaneous activity. There was no significant difference in network excitability between APV treated cultures and controls (see Figure 3B; *p* = 0.11), and network excitability in APV treated cultures did not significantly change during experiments (*p* = 0.13). In the control group network excitability slightly decreased with time (*p* = 0.045).

**Fig 3.**
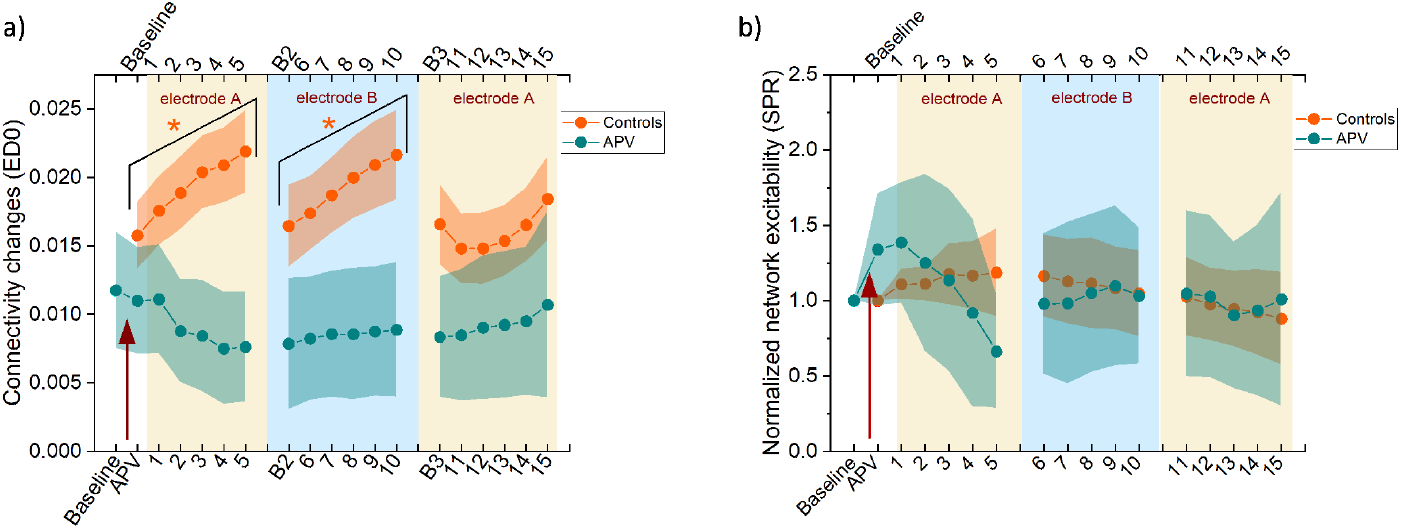
Memory formation and time course of network excitability. (a) shows stimulation-induced connectivity changes for controls (orange) and APV treated (green) cultures. First series of stimulation at electrode A induced connectivity changes in controls (*p* = 0.002) but not in APV experiments (*p* = 0.12). Stimulation at electrode B yielded similar effects (control *p* = 0.003, APV *p* = 0.88). Return to electrode A did not induce connectivity changes in either group (control *p* = 0.08, APV *p* = 0.61). (b) shows that network excitability was not affected by APV treatment (*p* = 0.11). In addition time was not affecting either group (control *p* = 0.13, APV *p* = 0.045). In both panels the red arrow indicates the addition of APV. Shaded areas show SEM, and indicate differences between experiments.

**Fig 4.**
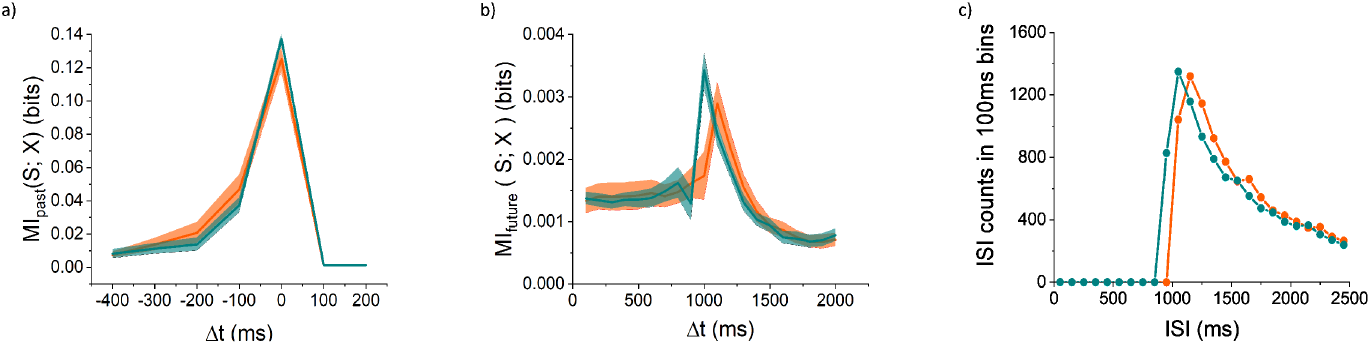
MI curves obtained in control (orange) and APV experiments (green). (a) *MI*_past_ shows a peak at Δ*t* = *−*200 *−* 0 ms corresponding to recorded stimulus responses. (b) *MI*_future_ shows a peak at Δ*t* = 1100 ms for control experiments and at Δ*t* = 1000 ms for APV experiments. Both follow the distribution of interstimulus intervals (ISIs) used and shown in (C). Shaded areas represents SEM.

### Memory trace formation

We first computed baseline functional connectivity fluctuations (*ED*_Baseline_) and possible drift (slope). in the networks. Baseline slopes averaged 3.6 × 10^−5^ ± 4.1 × 10^−5^ for 1^*st*^ stimulation of electrode A, −2.7 × 10^−5^ ± 2.3 × 10^−5^ for electrode B, and −3.2 × 10^−5^ ± 3.3 × 10^−5^ for return to electrode A (Figure 2). Figure 3 shows *ED*_Baseline_, which was used to compare stimulation-induced connectivity changes to spontaneous fluctuations in baseline.

Then we quantified connectivity changes induced by 10 min periods of low frequency stimulation at one of the electrodes (Eq. S2 Supplementary Information). Figure 3A shows that in the control group the first sets of stimuli at electrode A and the first sets of stimuli at electrode B induced significant connectivity changes (respectively *p* = 0.002 and *p* = 0.003). Interestingly, second sets of stimuli to electrode A did not induce further significant connectivity changes in the network (repeated measurement Friedman test: *p* = 0.08). In the APV group, stimulation at either electrode did not lead to significant connectivity changes (A first stimulation set *p* = 0.12, B stimulation *p* = 0.88, A second stimulation set *p* = 0.61).

### Quantification of prediction

We computed *MI*_past_ (Eq 1) and *MI*_future_ (Eq 2) to quantify short term memory and prediction. Figures 4A and B show *MI*_past_ and *MI*_future_ for control experiments (N=8) and APV experiments (N=10). Data from the control group has been used in an earlier study [34]. Both *MI*_past_ curves show a clear peak that corresponds to induced stimulus responses, which lasted up to *≈* 300 ms (Figure 4A). Both *MI*_future_ curves show an initial plateau until Δ*t* = 1000 ms, followed by a peak between Δ*t* = 1000 ms and Δ*t* = 1100 ms. These correspond to the most probable inter stimulus interval (see ISI distribution figure 4C). These peaks were followed by a decrease in *MI*_future_ for 1100 < Δ*t* < 2000 ms, reflecting the probability distribution of ISIs. The narrow shaded areas, that indicate SEM and represent differences between different hours of analysis across experiments, show that curves hardly changed during the 10 analyzed hours.

### Relationship between prediction and memory

Figure 5 shows how prediction, as quantified by Σ*MI*_future_ (Eq. S3 Supplementary Information), depended on short-term memory, as assessed by Σ*MI*_past_ (Eq. S4 Supplementary Information). Prediction was strongly correlated with short-term memory in control (*R* = 0.8 ± 0.07) as well as APV experiments (*R* = 0.87 *±* 0.09). Under control conditions the slope of this relationship decreased with time (*p* = 0.02) and the offset increased (*p* = 0.004). In APV experiments slope and offset did not significantly change in the considered 10h of stimulation (*p* = 0.56 and *p* = 0.79). See Figures 5C and D.

**Fig 5.**
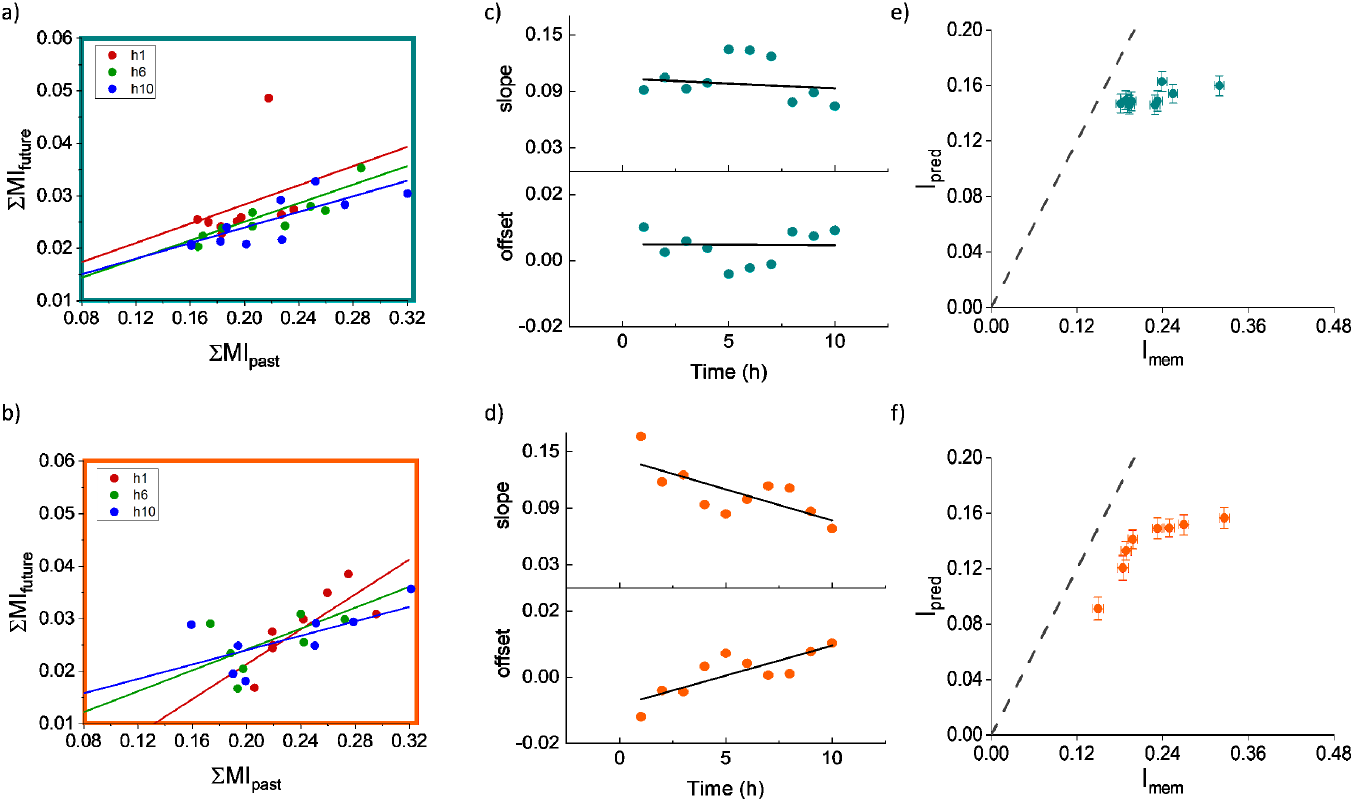
Relationship between prediction (Σ*MI*_future_) and short term memory (Σ*MI*_past_), and efficiency of prediction. A,B) examples (hour 1, 6 and 10) of the relationship between Σ*MI*_future_ and Σ*MI*_past_ and fitted trend lines in APV-treated cultures (a) and controls (b). (c) time course of slope and offset of fitted lines in (a). In APV experiments slope and offset did not significantly change with time (*p* = 0.82 and *p* = 0.97). (d) Time course of slope and offset of fitted lines in (b). In control experiments slope significantly decreased (*p* = 0.02) and offset significantly increased (*p* = 0.004) with time. (e) and (f) Efficiency of prediction in APV (e) and control (f) cultures. *I*_pred_ and *I*_mem_ were calculated using Eq 3, dashed lines indicate the theoretical optimum as derived in Eq. 4. This shows us the minimum memory necessary to enable a certain prediction.

### Efficiency of predictions

To study how efficiently APV-treated cultures predict compared to controls, we computed *I*_mem_ and *I*_pred_ (Eq. 3) for each culture, using the first 10 h of stimulation. Under control conditions cultures reached a predictive power (*I*_pred_) that was on average 62.7 ± 7% of the theoretical maximum (*R*(*D*)), (Figure 5F). When treated with APV cultures reached 69.6 ± 9% (Figure 5E). APV treated cultures were as efficient as control ones (*p* = 0.1).

## Discussion

The current study confirms that *in-vitro* networks of dissociated cortical neurons form memory traces when focally stimulated as shown in earlier work [22, 23]. Blockade of NMDA receptors impeded the formation of memory traces, demonstrating the pivotal role of NMDA receptor activation in the formation of memory traces. Networks under NMDA blockade were still able to predict upcoming stimuli. However, whereas prediction became less dependent on short-term memory with the formation of long-term memory traces in control cultures, this dependency did not decrease under NMDA blockade.

NMDA receptor activation plays an important role in STDP. The current finding that NMDA receptor blockade hampered long-term memory formation in cortical networks is in line with the notion that STDP is critically involved in this process. However, blocking a subset of excitatory receptors may also reduce network excitability, which has been reported to hamper memory formation [23]. We therefore verified that network excitability did not decrease significantly (Figure 3), which suggests that other mechanisms than reduced network excitability lead to impeded memory trace formation. Although NMDA block has been shown to affect the dynamics of network bursts [57], it did not impede network responses to electrical stimulation or the occurrence of spontaneous network bursts, which are the most plausible driving forces to reach a connectivity *<*=*>* activity equilibrium. Taken together, these observations are in agreement with a crucial role of STDP in memory trace formation [35, 45, 58, 59].

Information about future stimuli (*MI*_future_) in control and APV experiments were similarly shaped, with a plateau first, followed by a peak and subsequent decrease. *MI*_future_ is strongly influenced by the distribution of inter-stimulus intervals and the corresponding intrinsic predictive information of the stimulus vector about itself, as explained in earlier work [34]. The obtained curves suggest that upon receiving a stimulus at time t, the network infers the absence of other stimuli within the subsequent 1000 ms interval (nearly constant MI curve for Δ*t* ≤ 1000 ms, Figure 4). Furthermore, the network anticipates that the next stimulus is most likely to occur after 1000 *−* 1100 ms (peaks of MI curves), with a diminishing probability for longer intervals.

Prediction strongly depended on the presence of clearly recognizable stimulus responses. These stimulus responses are often interpreted as short-term memory of a stimulus, which has been suggested to depend mainly on sustained spiking in recurrent networks, creating attractor states [60]. Blocking NMDA receptors resulted in increasing stimulus responses during the first 5 hours after APV administration, followed by a decrease towards baseline values during the next hours (Figure S2). This suggests increasing excitability during the first 5 hours, which may reflect homeostatic mechanisms aiming to compensate for the blockade of NMDA receptors. On average, the NMDA antagonist did not reduce network excitability throughout the 10h analysis period. Networks remained responsive to focal electrical stimulation and they were still able to predict (Figure 4). This is in agreement with the notion that prediction, at least initially, depends on short-term memory, which is not blocked by the NMDA antagonist.

The slightly higher values of *MI*_future_ and the slightly higher prediction efficiency (Figure 4B and Figure 5E,F) might be due to the reduction of spontaneous activity and network bursts in APV-treated cultures, which yielded more recognizable stimulus responses in the recorded patterns (Figure 1C and D). Whereas prediction initially seemed to fully depend on short-term memory, this dependency decreased with time in focally stimulated networks under control conditions [34]. This type of stimulation induces long term memory traces in the network, suggesting that long-term memory traces are also involved in prediction under control conditions. This confirms earlier theoretical work on the role of long-term memory in prediction [61]. Interestingly, even though APV-treated networks were still able to predict, short-term memory dependency of prediction did not decrease with time (Figure 5D). Persistent short-term memory dependency was also observed in globally stimulated networks, where no long-term memory traces were formed [34]. This further supports the putative role of long-term memory in prediction.

Recent work suggested that unexpected stimuli contribute more to memory than expected ones [62–65]. Thus, there might be a relationship between the predictability of given stimuli and memory formation. Current results don’t provide insight because the timing and location of stimulation in memory experiments was deterministic, but this would be interesting to explore in future research.

The current study had a few limitations. The definition of memory differs from most other studies. This had the advantage that we were able to observe the formation of memory traces independently from recall. The mechanism of memory recall in our model is not clear yet, but the findings that stimulus responses stabilize during memory experiments, and start to appear in spontaneous firing patterns [22] support the concept of pattern completion [66, 67]. Further understanding of recall, preferably a quantitative measure of memory recall, could further confirm memory trace formation. It is not well-known how long the NMDA antagonist APV remains functional, and we could not directly estimate the half life of APV in the culture medium. However, indirect measures suggest that APV remained functional throughout experiments. No long-term memory traces were formed, even *>* 10 h after APV administration. Network excitability could not be measured in the prediction experimental protocol due to ongoing stimulation. However, the period of analysis in this protocol was limited to 10 h, and it seems unlikely that the effects of APV wore off more rapidly in this protocol than in the memory protocol. It would be interesting to see whether network excitability was affected by APV under continuous stimulation, as used in prediction experiments, but the method used to estimate network connectivity [55] requires longer periods of spontaneous activity between stimuli, containing a sufficient number of action potentials for statistical analysis. Controls and APV experiments were not performed on the same cultures, which may have introduced a bias related to possible culture differences.

In conclusion, We developed an in-vitro model that allows us to investigate cellular mechanisms that underlie memory and prediction. Blocking NMDA receptors in this model impedes long-term memory formation. Short-term memory of stimuli persisted under NMDA block, and networks were still able to predict. However, in contrast to control conditions, the dependency on short-term memory persisted throughout time. This further sustains the hypothesis that long-term memory becomes involved in prediction after extended exposure to a focal stimulus.

## Data availability

Part of the data (prediction control experiments) are available on Dryad (doi:10.5061/dryad.18931zd2t). All other data will be made available on Dryad upon acceptance.

## Acknowledgments

The authors would like to thank Gerco Hassink and Marloes Levers for the technical assistance, and Stephanie Palmer and Chris Hillar for valuable comments. This study was supported by the US Air Force Office for Scientific Research, Grant Number FA9550-19-1-0411.

## Author contributions statement

J.lF, S.M and M.L. conceived the study design, M.L. performed the experiments, M.L. J.lF. and analysed the data with the contribution of S.M., M.L. J. lF. developed the analysis scripts with a contribution from S.M., M.L. M.vP. J.lF. and S.M. wrote the manuscript draft. All authors reviewed the manuscript.

## Supplementary Information

### Additional analysis

#### APV concentration

We blocked NMDA receptors aiming to impede synaptic plasticity induced by NMDA receptor activation. Blocking a subgroup of excitatory receptors may also affect network excitability, which has been suggested as a possible determinant for memory functioning [23], and might thus bias results. We tested three APV concentrations: 15, 20 and 25 µM, in 5 cultures each. We first acquired 1 h of spontaneous activity (*Baseline*), and after APV administration we acquired an other hour of spontaneous activity (*Baseline*_*AP V*_) (see Figure S1). Periods of spontaneous activity (*Baseline* and *Baseline*_*AP V*_) were used to determine Mean firing rate (MFR), burstiness index (BI), and mean connectivity strength, which are related to network excitability. The MFR was computed as defined by Bologna et al and following Dias et al [23, 68]. BI was used to quantify the synchronicity of each culture following Wagenaar et al. and Dias et al. [23, 54]. Mean connectivity strength was determined applying CFP [20]. We computed the ratio (baselineAPV/baseline) of MFR, BI, and mean strength of functional connections to quantify the effects of APV on these network parameters.

### Responsiveness to stimulation

All memory experiments were performed using a constant stimulation frequency of 0.2 Hz, while in all prediction experiments inter-stimulus intervals were taken from a distribution with average stimulation frequency around 0.2 Hz. We chose this relatively low frequency to facilitate long lasting effectiveness of stimulation. We computed post stimulus time histograms (PSTHs) to assess the responsiveness of cultures to stimulation throughout the entire experimental protocol (memory: 17.5 h; prediction: 20 h). PSTHs were generated by summing all recorded activity in 5 ms bins from 300 ms before the stimulus until 500 ms after. In memory experiments PSTHs were computed for all 10 min stimulation periods, and in prediction experiments PSTHs were averaged per hour. To quantify network responsiveness to stimulation, we calculated the area under the curve (AUC), starting 15 ms after the stimulus (thus excluding stimulation artifacts and directly induced activity) and ending 500 ms after the stimulus. In memory experiments AUCs were normalized to the first 10 minutes of stimulation on that specific electrode, and in prediction experiments AUCs were normalized to the first hour. The duration of stimulus responses was determined from PSTHs as the maximum time after the stimulus where the average PSTH was still higher than the averaged activity before stimulation plus 5 times its standard deviation.

### Memory formation

To investigate memory formation we estimated changes in functional connectivity of the network. Functional connectivity was inferred from spontaneous activity recordings using conditional firing probability (CFP) [20]. This allows us to estimate strengths of connections between electrodes (groups of neurons), using the following equation:

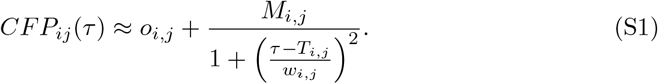

Here *M*_*i,j*_ is interpreted as the strength of the connection, *T*_*i,j*_ as the latency, *o*_*i,j*_ represents uncorrelated background activity, and *w*_*i,j*_ accounts for the width of the peak. For this analysis data were divided into chunks containing 2^13^ action potentials. This yielded a 60*x*60 connectivity matrix *M* for each chunk, that contained the strengths of connections between all possible pairs of electrodes. Formation of memory traces was investigated quantifying connectivity changes induced by each stimulation period by computation of Euclidean distances (EDs) between connectivity matrices *M* [23] using

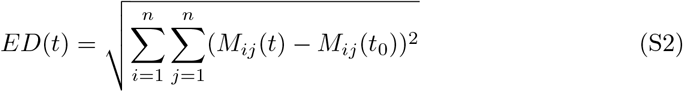

with *t* > *t*_0_.

### Relationship between memory and prediction

Estimates on the relationship between short-term memory and prediction were studied by computing the area under the curve of *MI*_future_ (Σ*MI*_future_) and *MI*_past_ (Σ*MI*_future_) for each hour of interest, for each experiment, using the following equations [34]:

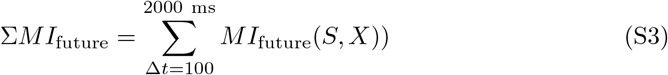

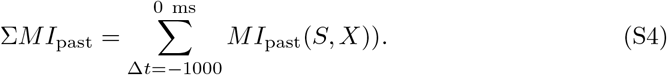

Here *S* indicates the binarized stimulation vector, and *X* refers to the binarized neuronal activity.

### Efficiency of predictions

To estimate predictions efficiency we follow the prediction bottleneck theory as explained in [34]. In short we want to understand which is the minimum memory to achieve a certain prediction. In a highly precise mathematical context, it emerges that retaining information concerning the minimal sufficient statistics of prediction, known as the forward-time causal states (*S*^+^), is crucial. Determining the essential components of these causal states necessitates numerical computation. For clarity, let’s designate 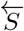 as the past and 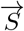 as the future of the input. Lets consider also 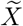 as representative of neuronal activity, then we wold like to have:

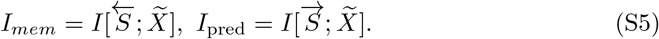

The predictive information bottleneck principle posits the simultaneous minimization of *I*_mem_ and maximization of *I*_pred_. It also states that with no memory (*I*_mem_ set to 0) there is no prediction (*I*_pred_ will also be 0). Conversely, if *I*_pred_ encompasses all information from the past to the future 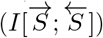, *I*_mem_ may become infinite due to the potential need to retain the entire past. Thus, the endeavor to reduce memory while enhancing predictive capacity entails a trade-off. However, the original formulation of the predictive information bottleneck faces a challenge: dealing with the entirety of the past and future presents mathematical impracticalities. Recent studies propose a solution: substituting the entire past of the stimulus with the time since the last stimulus (*S*^+^), and the entire future with the time to the next stimulus (*S*^−^), resulting in minimal information loss [56].

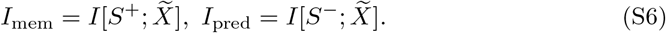

Memory embodies costs, while prediction signifies profit, yet memory remains indispensable for prediction. Rate-distortion theory, a facet of information theory, establishes the link between minimal resources and optimal precision, as described in [69]. In this way it is possible to estimate the trade-off between memory and prediction, as in equation 4 (main manuscript). The interpretation of this equation is that among all conditional probabilities (neural activity patterns) yielding predictive power of at least *D*, we seek the one requiring the least memory.

### Cultured neural network composition

We used cultures on coverslips with the same density as cultures on MEA to visualize the presence of both neurons and astrocytes under control conditions [53]. They were fixated after 19–26 days in vitro using 4% paraformaldahyde, permeabilized with 0.2% Triton X*−*100 and blocked with 1% BSA. Then they were stained for mouse anti-microtubuleassociated protein 2 (MAP2; M9942, 1:200; Sigma-Aldrich, St. Louis, MO) or goat anti-glial fibrillary acidic protein (GFAP; SAB250046, 1:200; Sigma-Aldrich) [53]. Secondary antibodies used were donkey anti mouse-CF555 (SAB4600060, 1:500; Sigma-Aldrich) and donkey anti goat-488 (SAB4600032, 1:500; Sigma-Aldrich). We also stained excitatory and inhibitory (pre) synapses. Fixation was the same as mention above but NGS was used as blocking buffer [52]. Primary antibodies used were: mouse anti vGLUT; Sigma-Aldrich (Cat. Nr. SAB5200262-100UG), and rabbit anti vGAT; Sigma-Aldrich (Cat. Nr. V5764-200UL). Secondary antibodies used were donkey anti mouse 555; Sigma–Aldrich (SAB4600060-250UL), goat anti mouse TR; Santa Cruz (sc-2983), and donkey anti rabbit FITC; Jackson ImmunoResearch (711-095-152). In addition, DAPI staining was applied(1:1000; Sigma-Aldrich, Cat. Nr. D9542-1MG) [52, 53]. All pictures were taken using a Nikon Eclipse 50i fluorescence microscope.

## Additional results

### APV concentration

We tested 3 different concentrations of APV (15, 20 and 25 µM). The averaged mean firing rate significantly decreased by about 50% for 15 (*p* = 0.026) and 20 µM of APV (*p* = 0.024), and remained unaffected by 25 µM (*p* = 0.34). The burstiness index tended to decrease (by *≈* 50%) for all concentrations, but changes were not significant (*p* = 0.19, *p* = 0.05 and *p* = 0.34, respectively). The mean connectivity strength tended to decrease for all concentrations, but changes were not significant(15 µM *p* = 0.08, 20 µM *p* = 0.34 and 25 µM *p* = 0.08). We chose to use 20 µM of APV because it showed the best trade-off between the effect on plasticity and reduction of network excitability.

### Responsiveness to stimulation

To assess responsiveness to stimulation throughout experiments, we calculated the area under the curve of post-stimulus time histograms as shown in Figure S2A and C. and the duration of stimulus responses as shown in Figure S2B and D. Two-way repeated measurement ANOVA showed that responsiveness did not significantly depend on time in the memory protocol (*p* = 0.29). Prediction data showed a significant interaction between time and APV application (*p <* 0.05), thus we performed one-way ANOVA to test the effect of time in controls and APV experiments separately. Stimulus response were not affected by time under control conditions (*p* = 0.68). However under APV administration responsiveness significantly depended on time (*p <* 0.001), with increasing responsiveness during the first 7 hours, followed by a decrease (Figure S2C). To avoid a possible bias of the quantification of prediction by the decreasing responsiveness, we chose to analyze only the first 10 hours. In both protocols, responsiveness did not significantly differ between conditions (control vs APV); memory protocol (*p* = 0.53), prediction protocol: *p* = 0.54), or on the two-way interaction between time and condition (*p* = 0.63). Figure S2B and D show that the duration of stimulus responses did not change with time (Memory protocol: *p* = 0.3; prediction protocol: *p* = 0.49), and did not differ between conditions (Memory protocol: *p* = 0.51; prediction protocol: *p* = 0.56), and there was no significant interaction (*p* = 0.71).

### Additional figures

**Fig S1.**
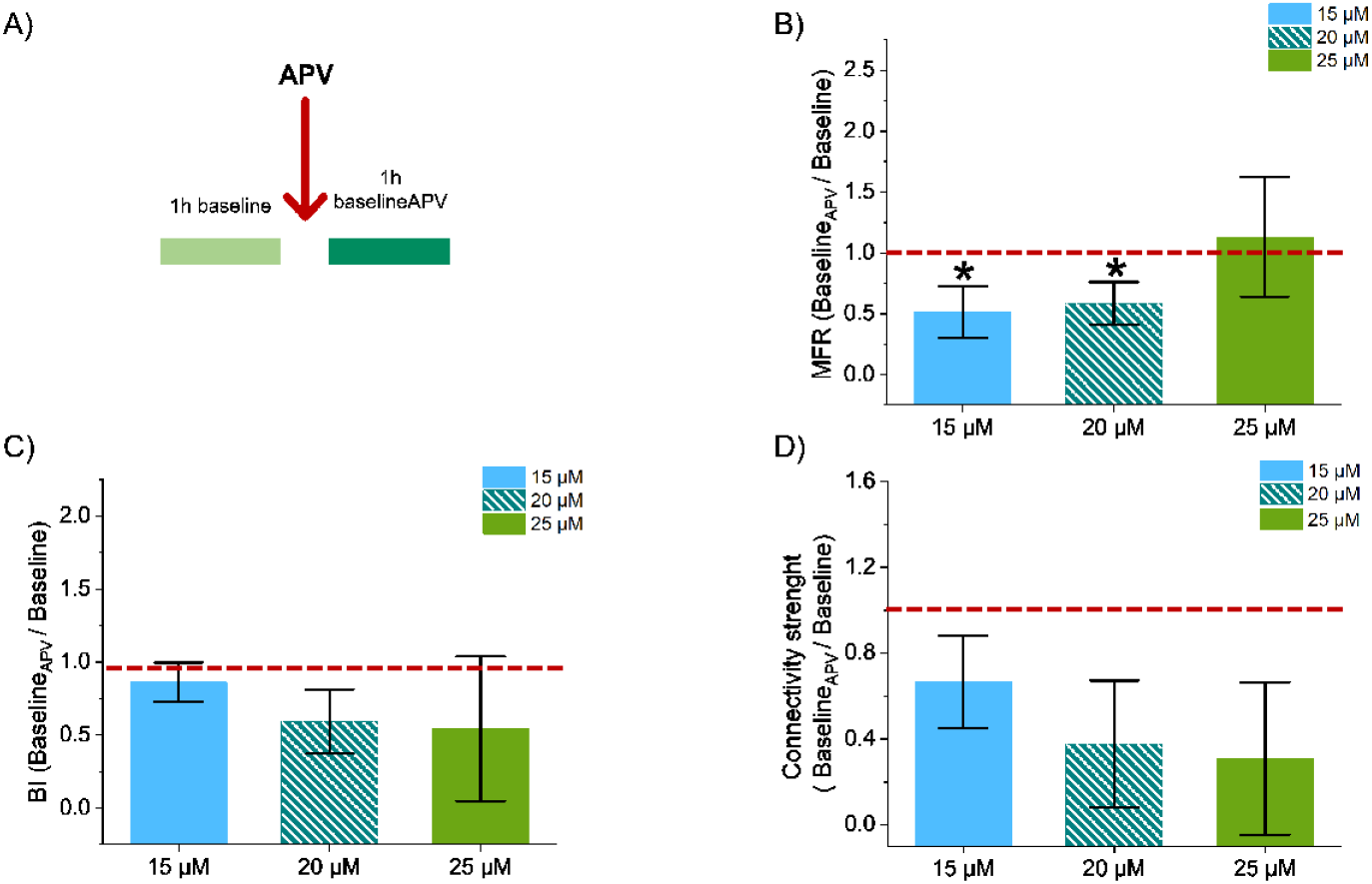
Effect of different APV concentrations on network excitability and plasticity. A) experimental protocol used to test each concentration. B) Effect of 15 (light blue), 20 (oblique stripes) and 25 µM (light green) of APV on mean firing rate (MFR). C) Effect of different APV concentrations on burstiness index (BI). D) Effect of the different APV concentrations on the mean connectivity strength. Significant changes are indicated by *.

**Fig S2.**
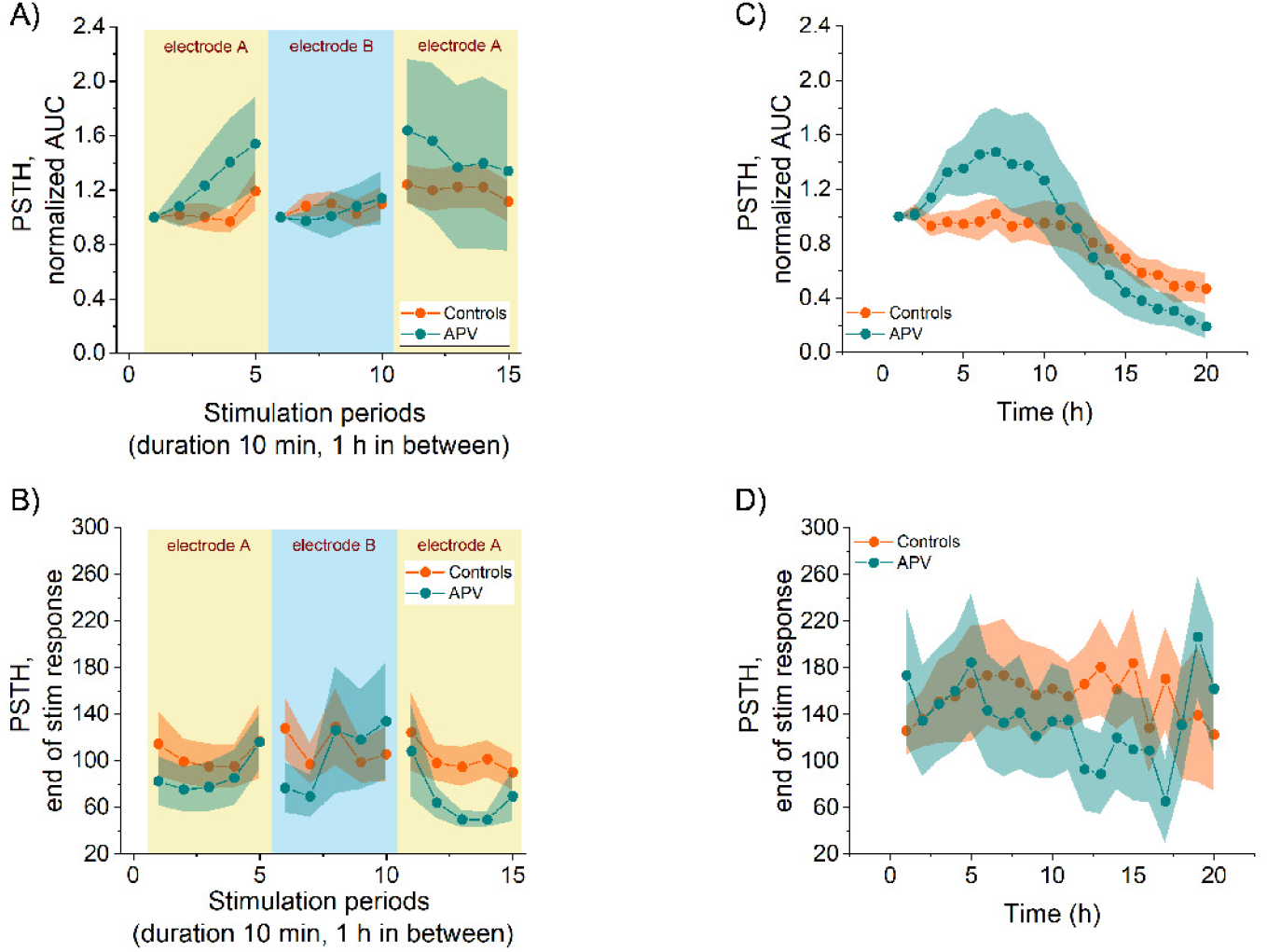
Effectiveness of stimulation. In all panels control experiments are shown in orange and APV ones in green. A) and B) are analysis related to the memory experiments. A) shows that there was no effect on the AUC of the PSTHs of either time (*p* = 0.29) or APV addition (*p* = 0.53). B) shows that there was no effect on the end of the PSTHs of either time or APV addition (respectively *p* = 0.3 and *p* = 0.51). C) and D) regard prediction experiments. C) shows that there was no effect on the AUC of the PSTHs induced by APV addition (*p* = 0.54), but in this case AUC was influenced by time (*p <* 0.001), while this did not happen under control conditions (*p* = 0.68). D) shows that there was no effect on the end of the PSTHs of either time or APV addition (*p* = 0.49 and *p* = 0.56 respectively). Shaded areas represent SEM.

**Fig S3.**
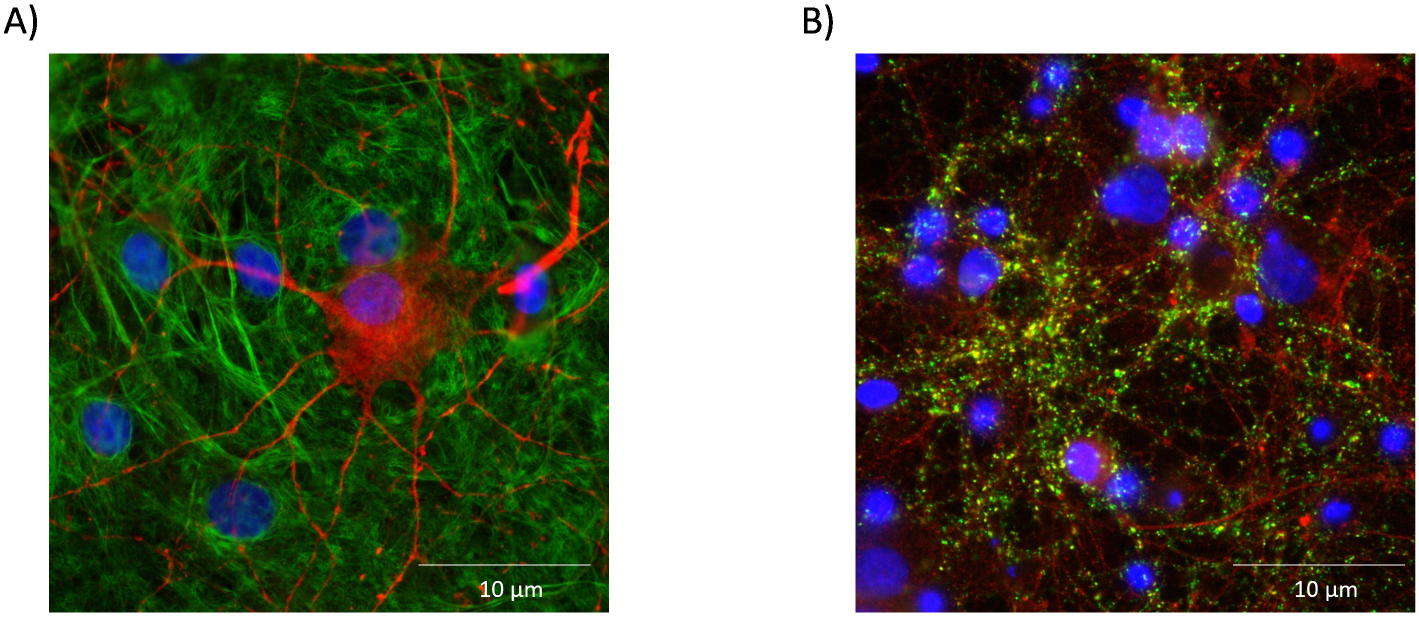
Cultured neural networks contained neurons and astrocytes, and excitatory and inhibitory synapses. A) Example of a culture, stained for MAP2 (neurons; red) and GFAP (astrocytes; green). B) Example of a culture stained for vGLUT (excitatory synapses; red) and vGAT (inhibitory synapses; green). In both panels cell nuclei were stained with DAPI (blue).

